# A deep learning model for fish classification base on DNA barcode

**DOI:** 10.1101/2021.02.15.431244

**Authors:** Lina Jin, Jiong Yu, Xiaoqian Yuan, Xusheng Du

## Abstract

Fish is one of the most extensive distributed organisms in the world, fish taxonomy is an important part of biodiversity and is also the basis of fishery resources management. However, the morphological characters are so subtle to identify and intact specimens are not available sometimes, making the research and application of morphological method laborious and time-consuming. DNA barcoding based on a fragment of the cytochrome c oxidase subunit I (COI) gene is a valuable molecular tool for species identification and biodiversity studies. In this paper, a novel deep learning classification approach that fuses Elastic Net-Stacked Autoencoder (EN-SAE) with Kernel Density Estimation (KDE), named ESK-model, is proposed bases on DNA barcode. In stage one, ESK-model preprocesses the original data from COI fragments. In stage two, EN-SAE is used to learn the deep features and obtain the outgroup score of each fish. In stage three, KDE is used to select the threshold base on the outgroup scores and classify fish from different families. The effectiveness and superiority of ESK-model have been validated by experiment on three dominant fish families and comparisons with state-of-the-art methods. Those findings confirm that the ESK-model can accurately classify fish from different family base on DNA barcode.

## Introduction

Fish is one of the most widely study group of aquatic organisms, about 27,683 fish species have most recently been catalogued into six classes, 62 orders and 540 families worldwide [1, 2]. Fish taxonomy and rapid species identification are the fundamental premise of fishery biodiversity and fishery resources management, and also an important part of marine biodiversity. As a traditional classification method, morphological identification has successfully described nearly one million species on the earth, which has laid a good foundation for species classification and identification [3, 4]. However, routine species classification poses a challenge for fish classification owing to four limitations. First, due to the differences of individual, gender and geographical, phenotypic plasticity and genetic variability used for fish discrimination can result in incorrect classification [5]. Second, with the deterioration of ecological environment and disturbance of human activities, many fishery resources have been seriously damaged, making it more difficult to collect fish specimens, especially for those with less natural resources [6, 7]. Third, some fishes show subtle dissimilarity in body shape, colors pattern, scale size and other external visible morphological features, which cause confusion of the same species. Finally, the use of key not only demands professional taxonomic knowledge, but also requires extensive experience that misdiagnoses are common [8]. The limitations of morphology-based method, a new technology to fish classification is needed.

Genomic approach is a new taxonomic technique combining molecular biology with bioinformatics that uses DNA sequences as ‘barcodes’ to differentiate organisms [5]. The DNA-based barcoding method is attainable to non-specialists. Many studies have shown the effectiveness of DNA barcode technology for more than 15 years, it has been extensive used in various fields such as species identification [9], discovery of new species or cryptic species [10, 11], phylogeny and molecular evolution [12], biodiversity survey and assessment [13, 14], customs inspection and quarantine [15], conservation biology [16].

In the field of species classification, a short gene segment is used in DNA barcoding, called the COI sequence, to build global standard dataset platforms, universal technical rules and identification systems for animals’ taxonomy [1]. COI gene has the characteristics of high evolution rate, obvious interspecific variation, relatively conservative within species, good universality of primers and easy amplification [17]. Therefore, COI gene has been widespread employed as an effective DNA barcode for species classification of varied animal lineages, including bird [18, 19], Mosquito [20, 21], marine fish [22–24], freshwater fish [25–27]. DNA barcode based on COI gene can be used to identify marine fish up to 98%, while freshwater fish can be identified with 93% accuracy [28]. The approach base on DNA barcode has been proven to be a valuable molecular tool for fish classification.

However, the complexity and high-dimensional characteristics in COI gene sequences, analyzing these sequences reasonably and obtaining accessible information that humans can classify fishes correctly are a major challenge. This issue requires a multidisciplinary approach to deal with DNA sequences and to analyze the information contained from data. Deep learning, a method of learning and extracting useful representations from raw data, trains model, and then, uses the model to make predictions, has made great progress in recent years [29]. Therefore, in this paper, we propose a novel approach based on DNA barcode, use the deep learning model to classify fish from different families and determine which fishes are regarded as outgroup, called ESK-model. To verify the effectiveness of the model, three families with many species and obvious interspecific variation were selected as the datasets. First, the model preprocesses the original data that makes the COI gene sequences into a matrix representation, then, converts them into numerical data. Second, the model learns these data using EN-SAE model and obtains an outgroup score of each fish. Finally, the KDE model is used to generate a threshold and to predict which fish is outgroup base on threshold. The main contributions of our paper are as follows:

- We introduce a deep learning model to classify fish from different families and determine which fish is outgroup based on DNA barcode, which is effective and robust.
- To solve the model overfitting caused by COI gene sample of species in the same family is limited, an Elastic Net is used for the model to increase the generalization ability.
- We employ EN-SAE model to receive outgroup scores. The decision threshold is automatically learned from organisms in same family by KDE model. An original predictor is proposed based on the anomaly scores, while other classification works often omit the importance of automatic learning threshold.
- We quantitatively evaluate the performance of our approach, and the results demonstrate that our ESK-model outperforms state-of-the-art methods.

## Materials and Methods

### Data description

The COI sequences from three dominant families of fish in this study were obtained from GenBank(www.ncbi.nlm.nih.gov), including Sciaenidae, Barbinae and Mugilidae. Among them, Sciaenidae and Mugilidae belong to marine fish, Barbinae belongs to freshwater fish. The genetic relationship and molecular divergence are considered for selecting outgroups. The relevant information concerning the features, specimen size and outgroup ratio of three families were summarized in Table 1.

- Sciaenidae. The COI fragments contained 307 individuals of 21 species, 13 genera in Sciaenidae family. 18 homologous sequences in *Nemipterus virgatus*, *Epinephelus awoara, Leiognathus equulus* and *Leiognathus ruconius* were selected from different families, which were under the same order as Sciaenidae. After processing, the length of COI gene fragment was 596 bp. Species of experimental samples on Sciaenidae is shown in S1 Table.
- Barbinae. A total of 998 individuals from 103 species pertaining to 9 genera of Barbinae were barcoded, which were 544 bp of COI gene sequence length. In addition, 24 homologous sequences from 6 genera including *Foa brachygramma* and *Cheilodipterus macrodon* belong to Apogonidae were used as outgroup. Species of experimental samples on Barbinae is shown in S2 Table.
- Mugilidae. In this dataset, 776 Mugilidae sequences from 23 species belong to 7 genera were collected, which the length of COI gene was 565 bp. 20 homologous sequences in *Sphyraena pinguis* and *Sphyraena jello* from Mugiliformes were designated as outgroup. Species of experimental samples on Mugilidae is shown in S3 Table.

**Table 1.** Summary of datasets.

| Family | Feature | Instance | Outgroup ratio (%) |
| --- | --- | --- | --- |
| Sciaenidae | 596 | 325 | 5.54 |
| Barbinae | 544 | 1022 | 2.35 |
| Mugilidae | 565 | 796 | 2.51 |

### Data preprocessing

#### Data definition

To facilitate the subsequent processing, DNA sequences can be represented by a matrix. The COI sequences for each family were formulated as follows:

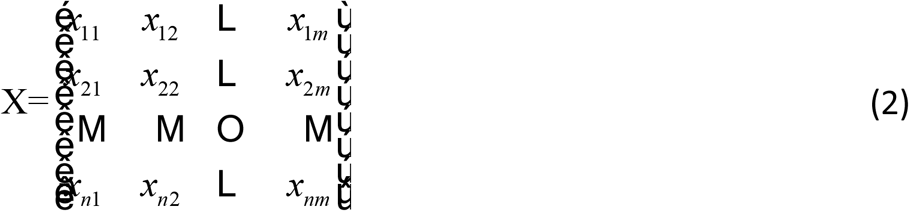

where n denotes the size of samples, and m denotes the number of features in each species.

#### One-hot code

One-hot encoding is the process of converting categorical variables into a form that is easy to use by machine learning algorithms, which are a combination of 0 and 1 [30]. Therefore, the model encodes matrix into a numeric type of data using one-hot code. COI gene is composed of four bases, A, T, C, G. Each coded base was a 1×4 vector [0, 0, a_i_, 0], where a_i_=1.Therefore, four bases were formulated as follows:

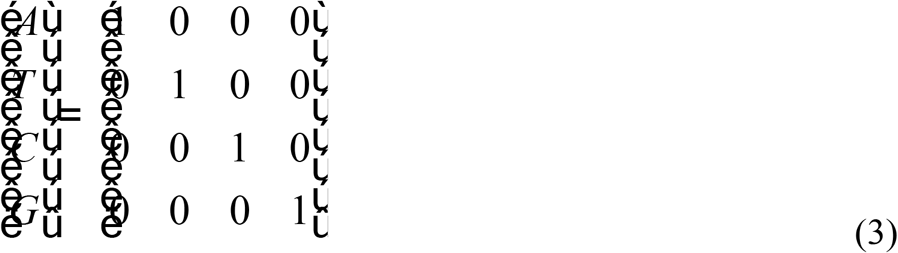

### Method introduction

#### An overview of the ESK-model

An overview of the proposed model is shown in Fig 1, ESK-model, which consists of three stages: (1) the data preprocessing stage, (2) learning deep features and computing each species outgroup score stage, and (3) deciding threshold base on outgroup scores and classifying fishes from different family stage.

**Fig 1.**
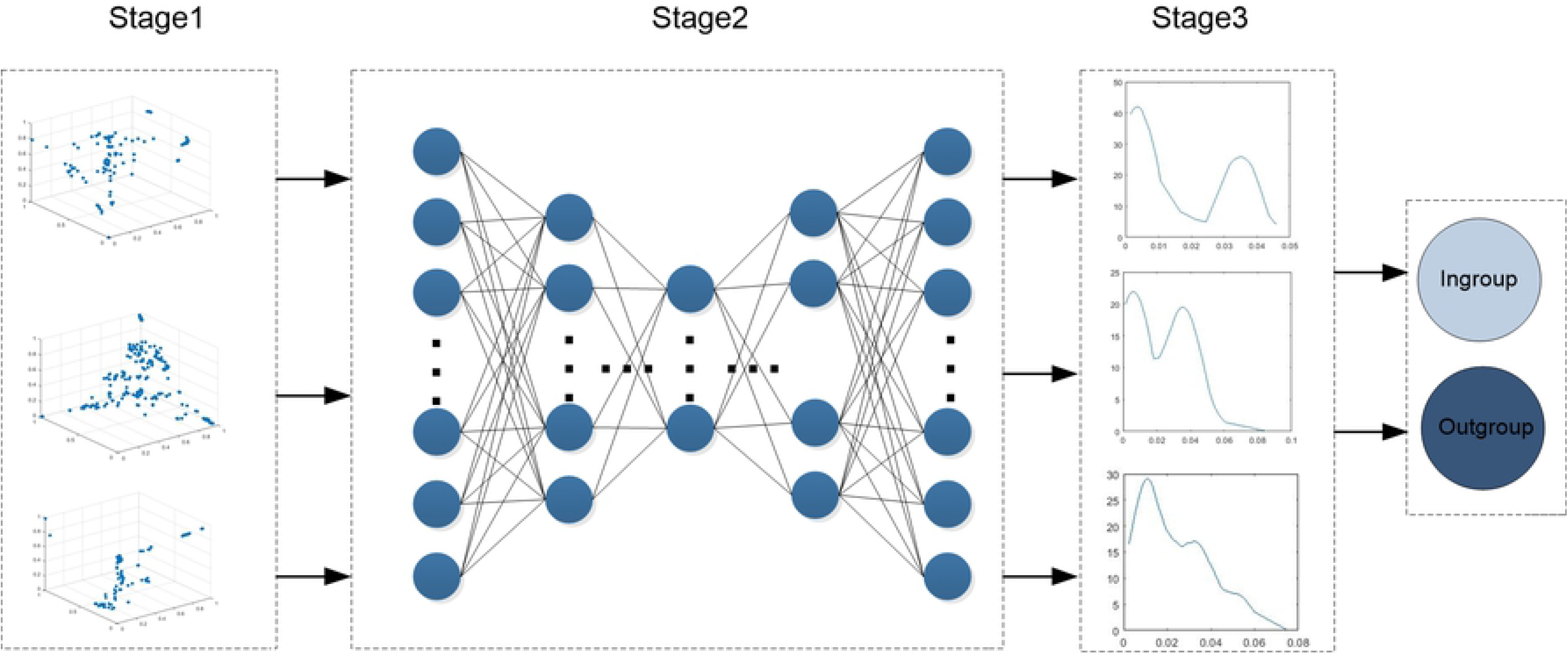
An overview of ESK-model. Three-dimensional visualization of data is shown in Stage1, the distribution of the anomaly scores is shown in Stage 3.

In stage one, there are two main tasks: (1) preprocessing raw data by representing the COI gene sequence in a matrix and (2) the one-hot code is performed on the matrix because the features of each fish species need to be transformed into numerical data. Finally, the preprocessed data are used as inputs for stage two.

In stage two, a deep learning network, EN-SAE, is used to learn deep features from the data preprocessed in stage one. The model utilizes the EN-SAE model to compress the digitalized data into a representation of the potential data to reconstruct input, then, calculates the difference between input and output, and obtains an outgroup score of each fish. Finally, the outgroup scores are used as inputs for stage three.

In stage three, the KDE technique is used to learn the relationship between each score from stage two, and then, fits the data distribution according to properties of the outgroup scores. After that, the KDE model determines which fish is inner group and which fish is outer group base on the threshold.

#### Learning deep features and computing outgroup scores by EN-SAE

Traditional AE is a three-layer neural network, including an input layer, an output layer and a hidden layer. The structure of AE is symmetric, that is, the input layer and output layer have the same number of nodes and the dimensions of each node are the same too [31]. The purpose of AE is to compress input data and save useful information to reconstruct input, and use the back propagation algorithm to update the weights so that the output data is as similar to the input data as possible [32]. However, the output data are not sufficient to yield a rewarding representation of input. The reconstruction criterion with three-layer structure is unable to guarantee the extraction of useful features as it can lead to the obvious solution “simply copy the input” [33]. The SAE can greatly solve this problem.

The SAE model builds a deep neural networks base on AE by stacking several AEs, puts the hidden representation of the upper layer as the input of the next AE. In other word, extracting the compressed features of hidden layer into next AE to training. In this way, training layer-by-layer can achieve input features compressed. At the same time, more meaningful features of COI sequences are obtained. The decoder can be reconstructed back into the input with a sufficiently small differences, the structure of SAE is expressed in Fig 2.

**Fig 2.**
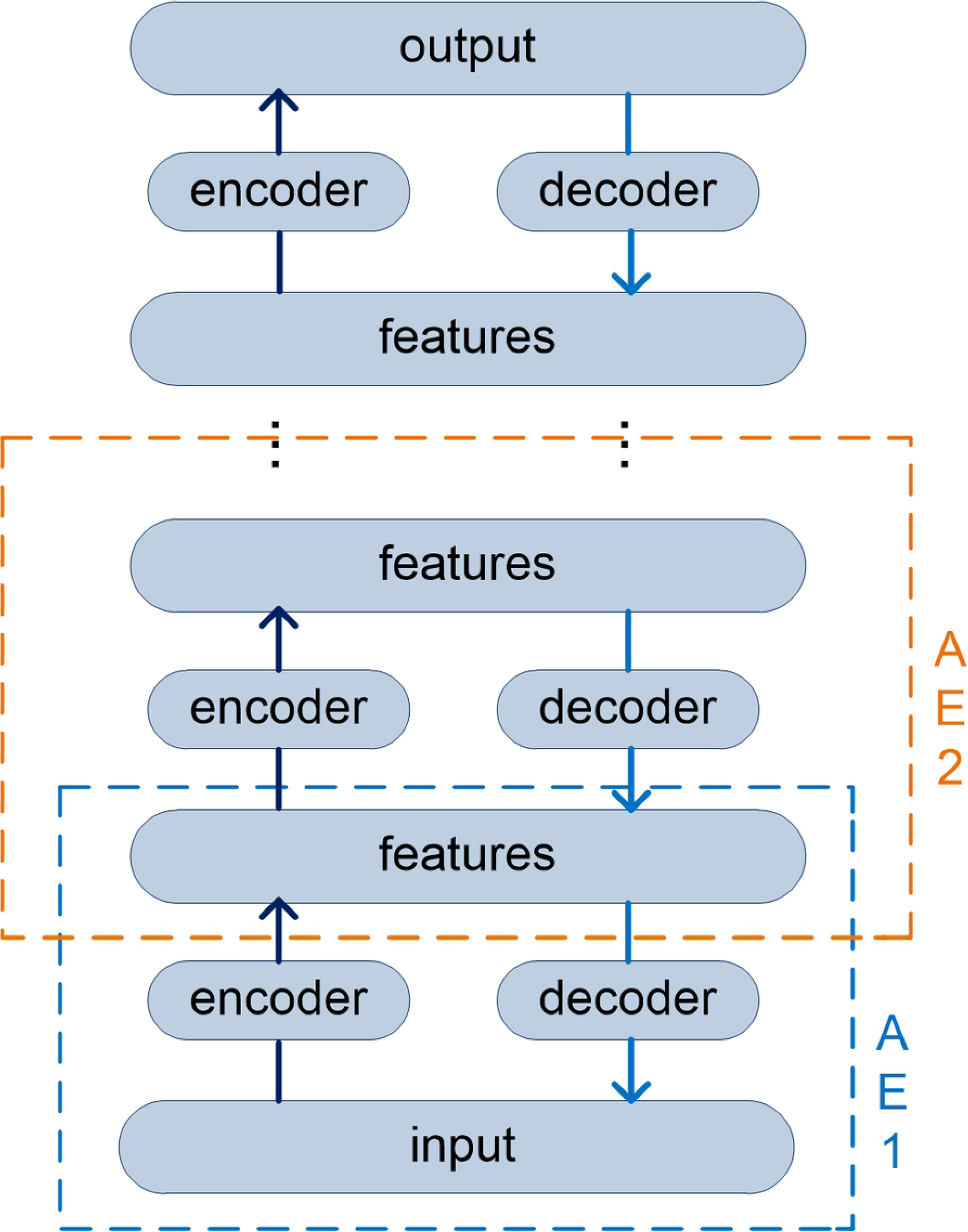
The structure of SAE.

There are two basic steps in SAE training: encoder and decoder.

1. Encoder: in this step, the activation function σ_e_ maps input data vector x to hidden representation h that can compress the input data and retain more useful representation, the typical form followed by a nonlinear representation:

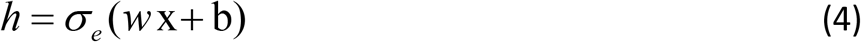

where x denotes input data vector, w is a weight matrix connecting the input layer to hidden layer, b is bias vector belongs to nodes of latent layer, σ_e_ represents activation function, such as Sigmoid, Relu, Tanh, etc.
2. Decoder: in this step, the hidden representation h is mapped into reconstruction vector y, the typical form as follows:

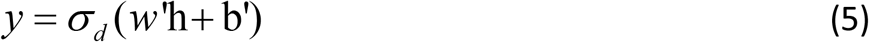

where w’ is weight matrix connecting the latent layer to output layer, b’ is bias vector, σ_d_ represents activation function.

Loss function is defined to measure the reliability of SAE. SAE is trained to reconstruct the features of input, and the weight of encoder and decoder are adjusted to minimize the error between output and input. Thus, loss function is introduced, it is represented by mean square error as follows:

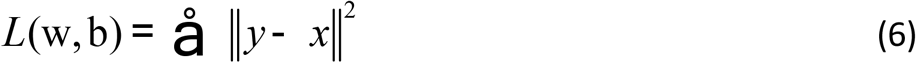

However, a COI gene fragment has too many features, which leads to the high dimensionality of the training data. At the same time, fish species contained in each family are limited, resulting in a relatively small dataset. Therefore, the model cannot fully learn the characteristics of each fish species. In order to improve the generalization ability of the proposed model, make the structure risky minimize, add some kinds of constraint, reduce the weight of useless features. Base on this point, Elastic Net composed of L1-norm and L2-norm is proposed in this method. The structure of EN-SAE model is shown in Fig 3. It can also treat L1-norm and L2-norm as penalty for loss function to restrict some parameters in the process of training.

**Fig 3.**
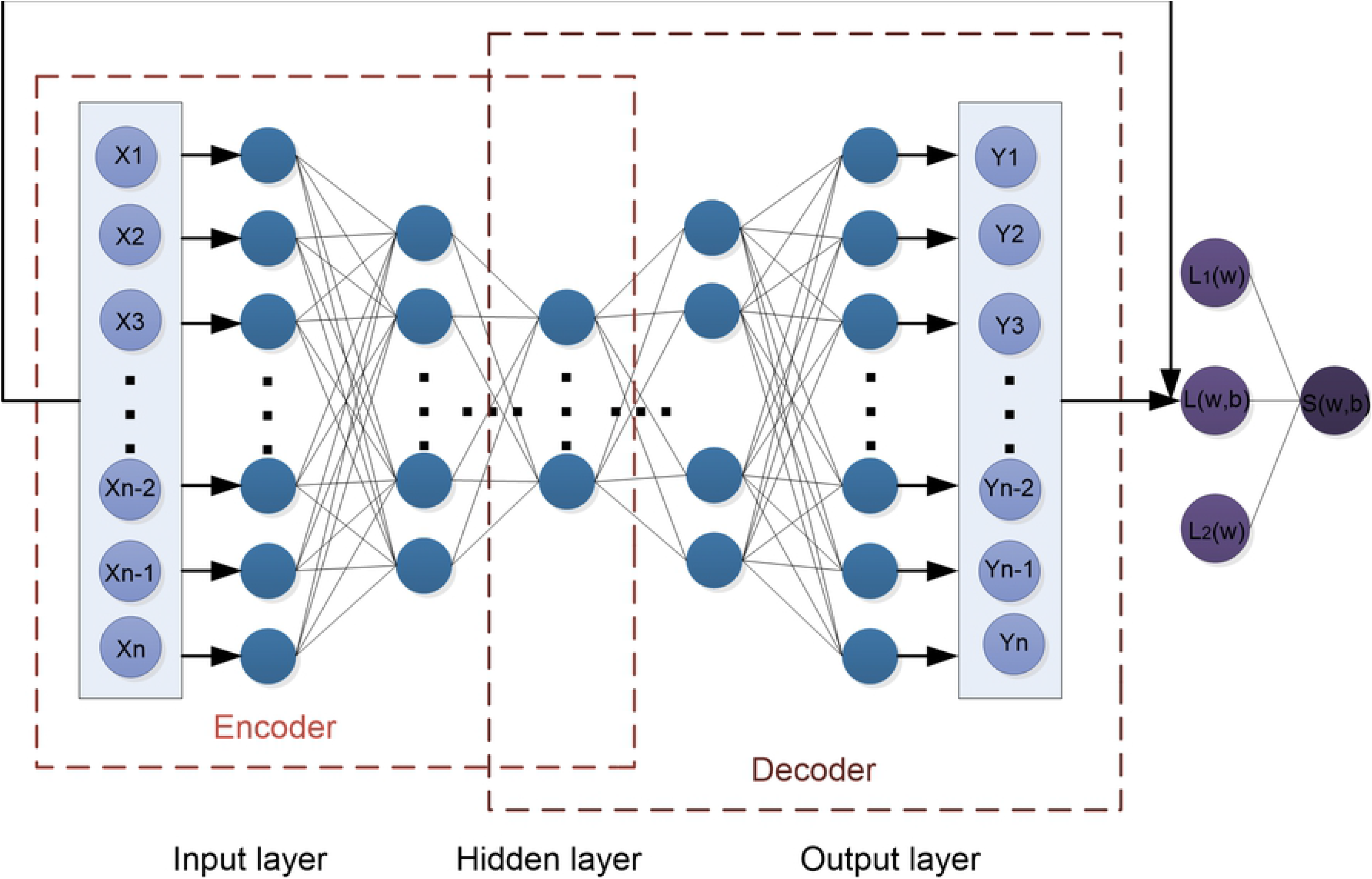
The structure of EN-SAE model.

L1-norm also called Lasso regression, which contributes to generating a sparse matrix. And it is defined as: 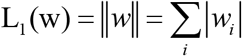, where is the sum of the absolute value of each element in weight vector w. Thus, it can be used to choose more meaningful representations. When training model, the features are too many to select what are contribute more for this model. So we dropped the connections that the contribution of this model is so tiny, even if drop its have no impact on the model [34]. It can reduce time consuming and study more useful features.

L2-norm also called Ridge regression, which is defined as: 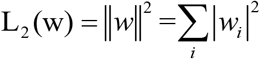, where is the sum of the squares of each element in weight vector w. In the process of training, we usually tend to make the weight as small as possible, because it is generally believed that the model with small parameters is simpler and can fit different data effectively. Thus, L2-norm can void overfitting to some extent and improve the generalization of model to adapt different fish families.

On the basis of proposed EN-SAE model, the outgroup score of each species can be defined to measure whether fish is outgroup. The higher outgroup scores are, the more likely they are to be treated as outgroup.

Therefore, the outgroup scores can be calculated by the following formula:

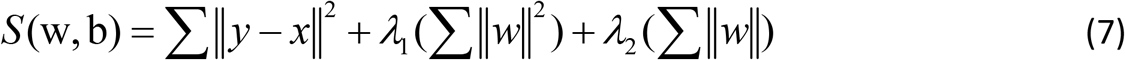

where λ_1_ is a parameter to adjust the L2-norm, λ_2_ is a parameter to adjust the L1-norm.

The EN-SAE model rejects high-dimensional features into low-dimensional features step by step to obtain higher representation of COI sequences, which is significantly more suitable for extract features and express data from original data.

#### Analyzing the outgroup scores by using KDE

KDE borrows its intuitive approach from the familiar histogram, which is among the most common nonparametric density estimation techniques. KDE provides a method of smoothing data points, and then, the distribution is fitted by the properties of data itself. The decision threshold is ascertained by using KDE model base on the outgroup scores. After that, the correct classification results of fish will be found. Given the outgroup scores vector s, which obtained from EN-SAE model, KDE estimates the probability density function (PDF) p(s) in a nonparametric way:

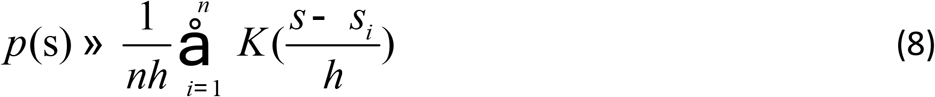

where n is the size of the training dataset, {s_i_}, i = 1, 2,…, n, is the training dataset’s outgroup scores vector, K (⋅) is the kernel function, and h is the bandwidth.

There are many kinds of kernel function, epanechnikov function is the most common function in density estimation and also has a good effect. Therefore, the epanechnikov is used to estimate the PDF:

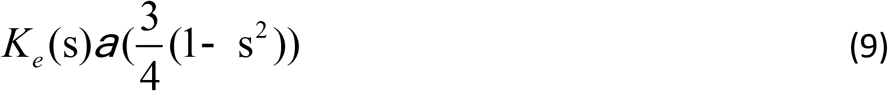

After obtaining p(s) of training the outgroup scores vector s by KDE, the cumulative distribution function (CDF) F(s) can be defined as fellow:

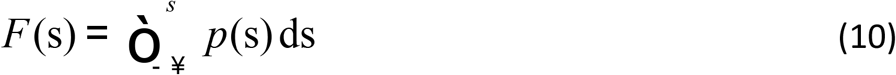

Given a significance level parameter α ∈ [0,1] and combine with CDF, a decision threshold s_α_ can be found, s_α_ satisfies following formula:

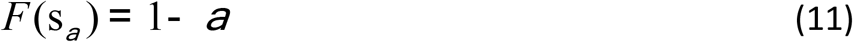

If the outgroup scores of each species meet the condition s ≥ s_α_, this species will be considered as outgroup. On the contrary, they are ingroup. Confirmed by repeated experiments that significance level parameter α is recommended to be set to 0.05. ESK-model algorithm is summarized as shown in Algorithm 1.

**Algorithm 1.**
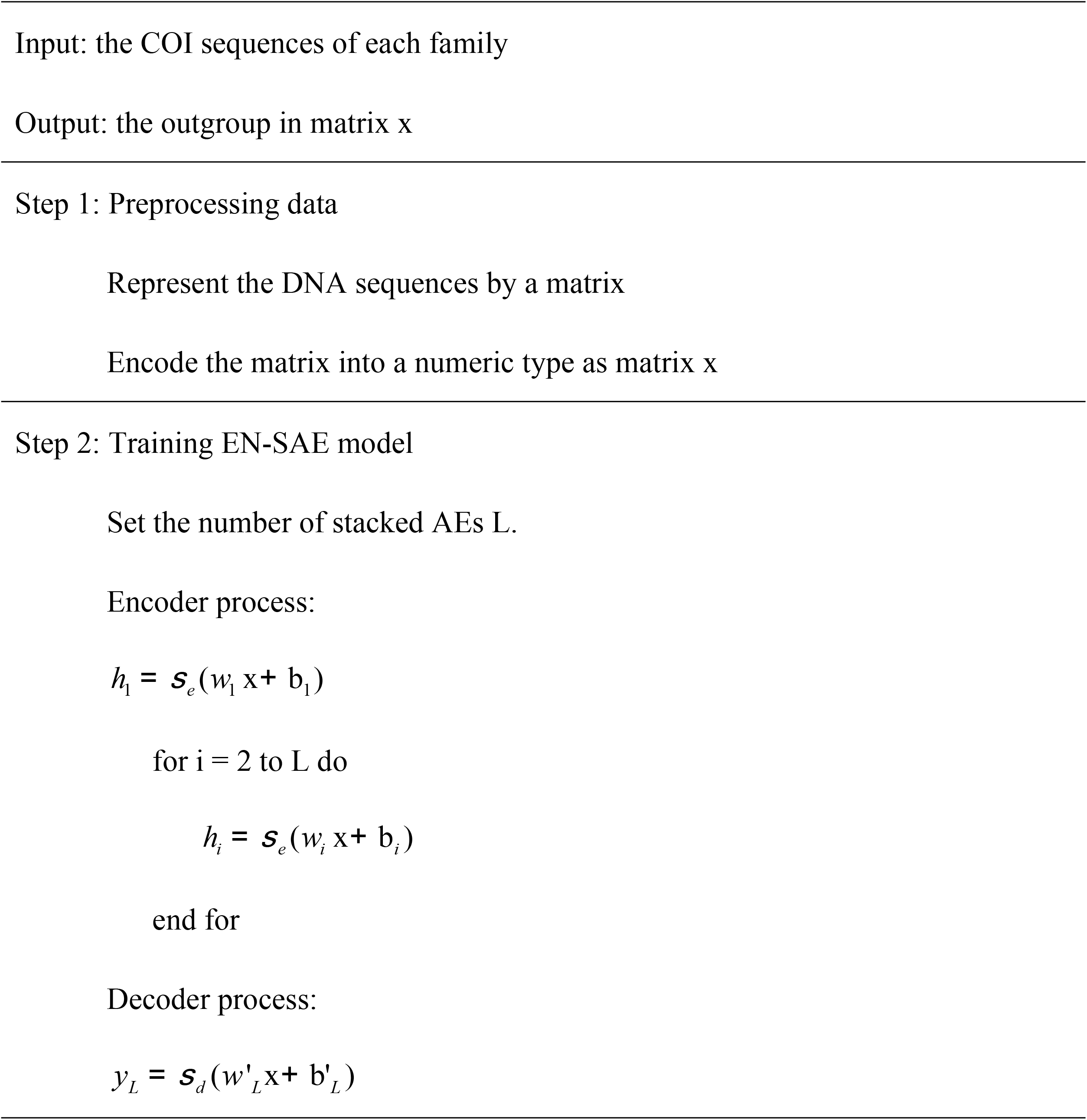
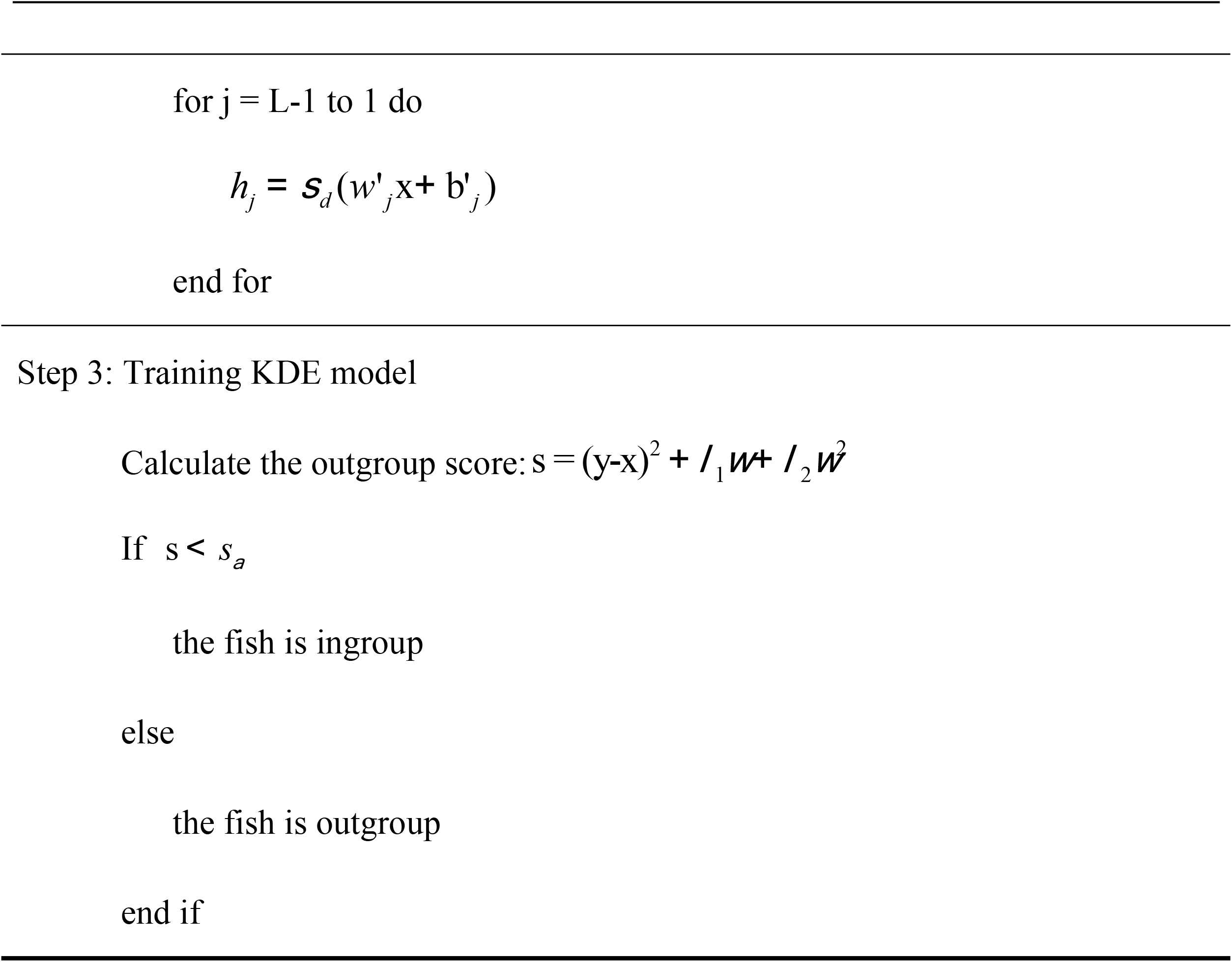
ESK-model

### Evaluation method

To test performance of the proposed model, divide the sample into four situations based on the actual classification and the ESK-model predicted classification. In Table 2, four situations are illustrated with a confusion matrix. True positive (TP) is the number of outgroups that are correctly classified as outgroup. True negative (TN) is the number of ingroups that are correctly classified as ingroup. False positive (FP) is the number of ingroups that are wrongly classified as outgroup. False negative (FN) is the number of outgroups that are wrongly classified as ingroup.

**Table 2.**
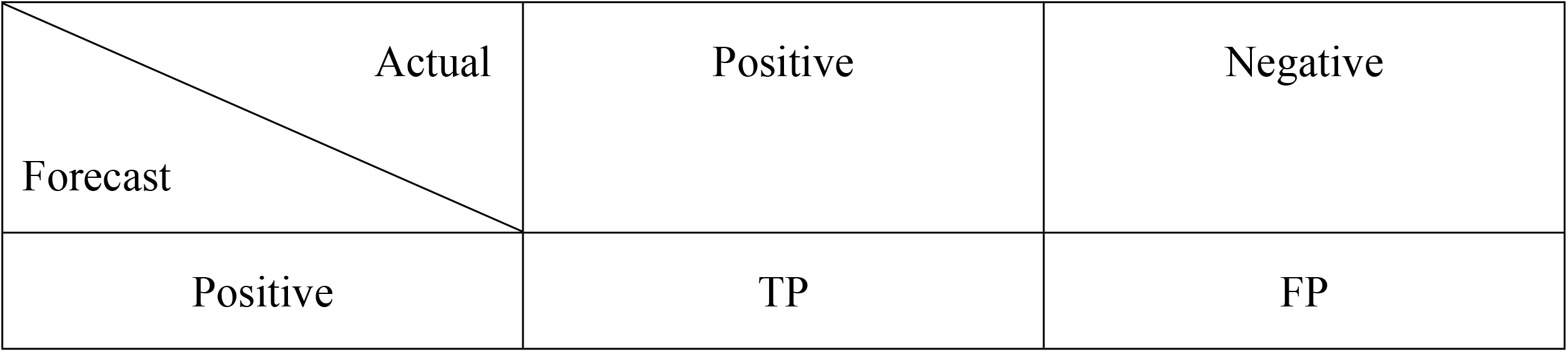
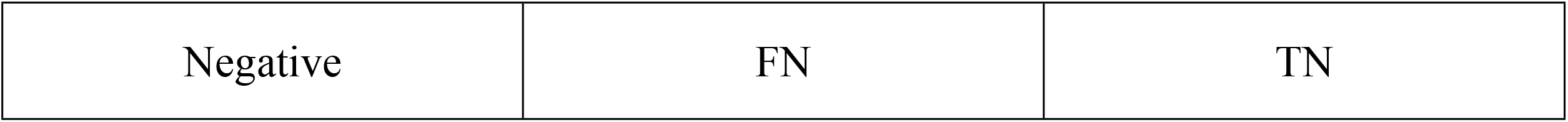
Confusion matrix.

With the confusion matrix, the classification performance of all experiments was measured by three criterions for Accuracy, Recall, and F-Measure. Those evaluation equations are formulated as follows:

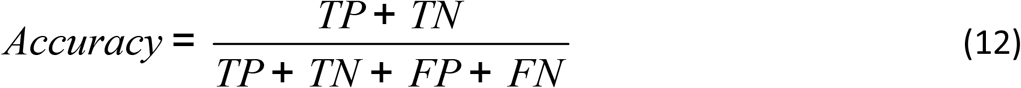

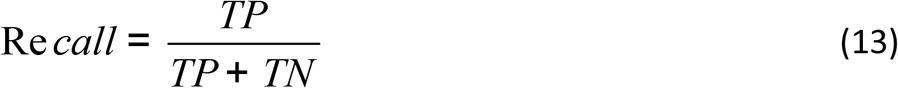

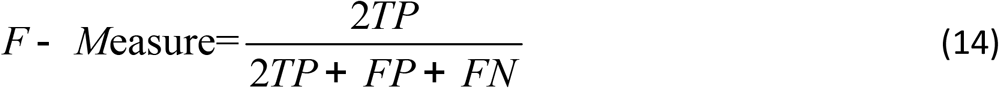

## Result

### Impact of the number of stacked AEs on the classification performance

In the field of deep learning, the number of layers in the model is a critical factor, because it directly affects the performance of the model. After all of the COI sequences were prepared, the impact of the number of stacked AEs in our model on the classification performance was also assessed. The outgroup scores trend with various stacked AEs from 3 to 8 on Sciaenidae, Barbinae, Mugilidae is shown in Figs 4-6. The experimental results showed in Fig 4 demonstrate that, as the number of AEs increased, the outgroup scores decreased rapidly on Sciaenidae when the number of AEs was fewer than five. The outgroup scores gradually stabilized when the number of AEs was greater than five. The outgroup scores on other two datasets showed the same trend as those on Sciaenidae. These results reach the best classification performance when the number of AEs was stacked to five.

**Fig 4.**
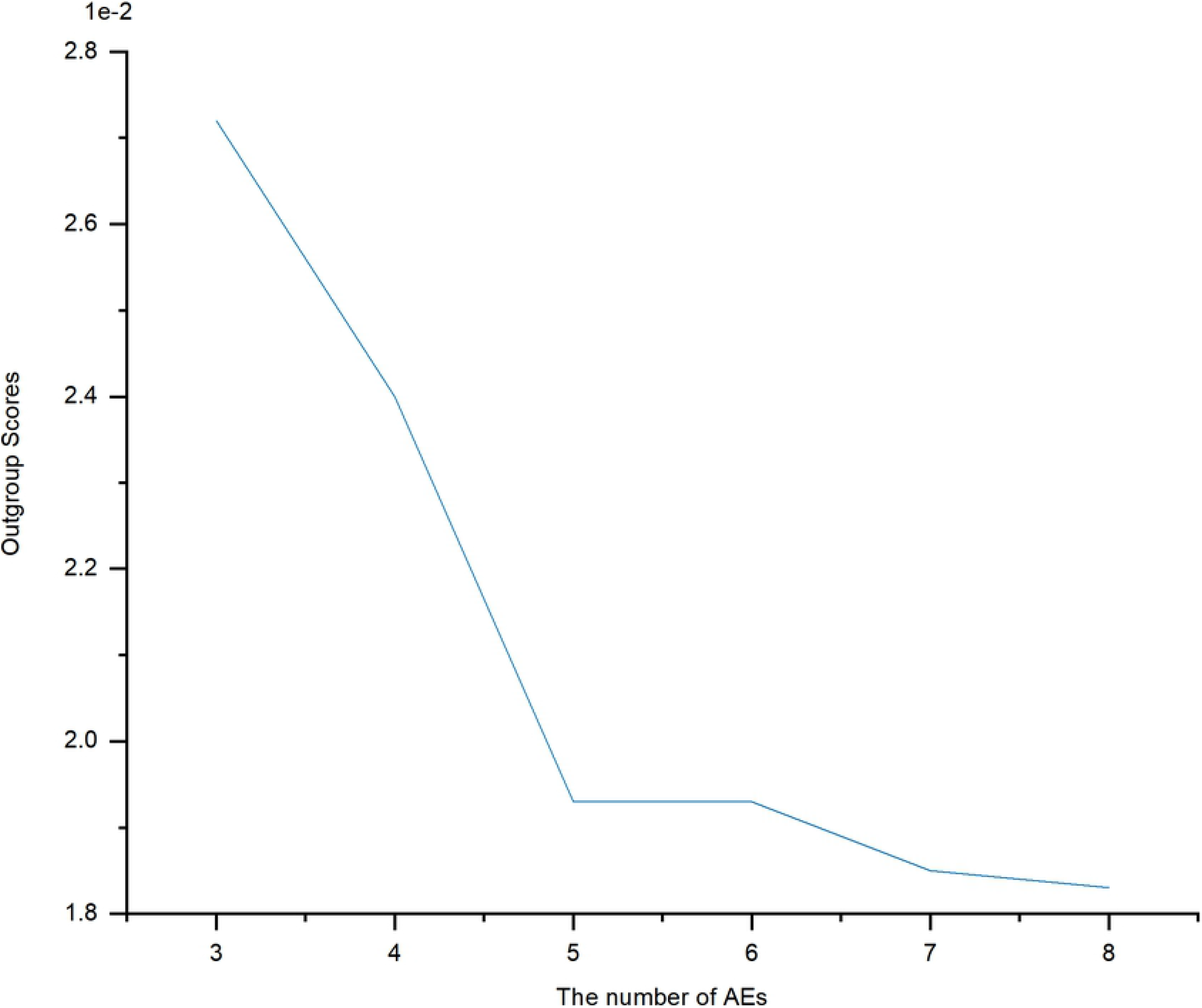
The outgroup scores trend on Sciaenidae.

**Fig 5.**
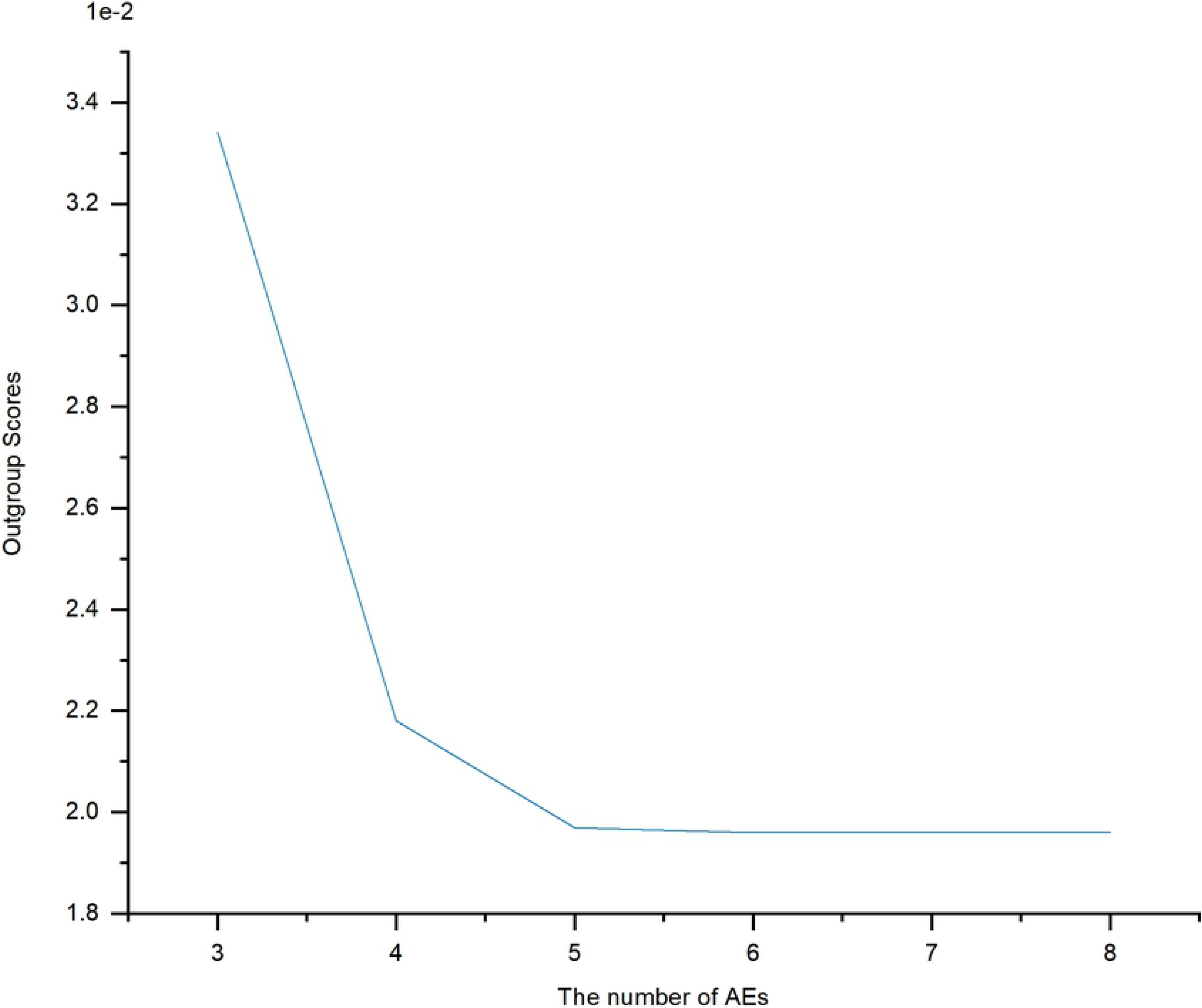
The outgroup scores trend on Barbinae.

**Fig 6.**
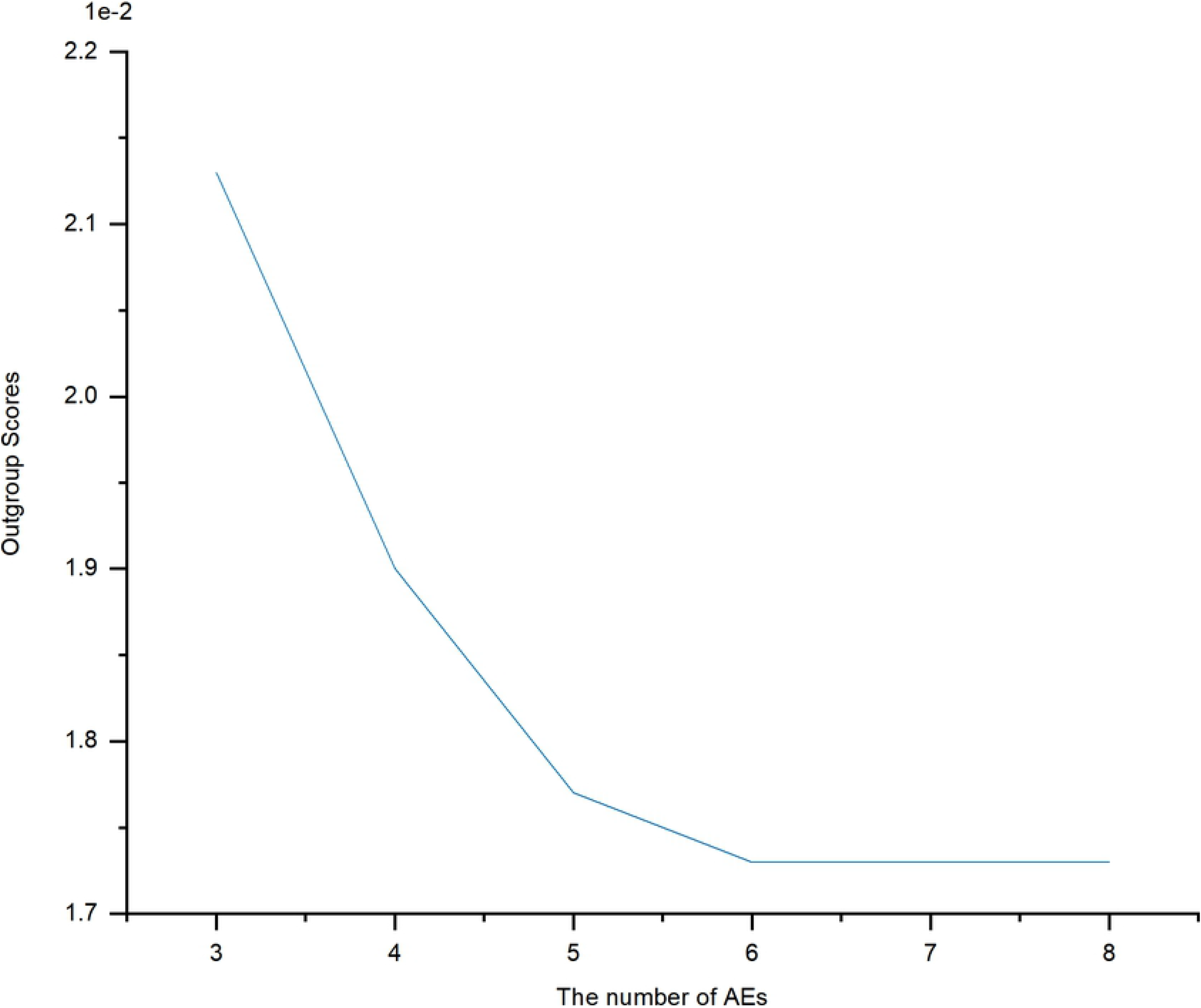
The outgroup scores trend on Mugilidae.

Additionally, Table 3 illustrates the detailed data corresponding to Figs 4-6. The results of Table 3 show that the outgroup scores of proposed model with five layers on different datasets were 0.0193, 0.0197 and 0.01, respectively. Moreover, after the number of AEs increased from 3 to 5, the outgroup scores on three datasets decreased by approximately 29.04%, 41.02% and 16.90%, respectively. Those results indicate that the proposed method can achieve low scores on identifying fish from different families and the outgroup scores tend to be stable gradually.

**Table 3.** The outgroup scores with different numbers of AEs on three datasets.

| Layers | Sciaenidae | Barbinae | Mugilidae |
| --- | --- | --- | --- |
| 3 | 0.0272 | 0.0334 | 0.0213 |
| 4 | 0.0240 | 0.0218 | 0.0190 |
| 5 | **0.0193** | **0.0197** | **0.0177** |
| 6 | 0.0193 | 0.0196 | 0.0173 |
| 7 | 0.0185 | 0.0196 | 0.0173 |
| 8 | 0.0183 | 0.0196 | 0.0173 |

Note that the bold values denote the outgroup scores with five stacked AEs.

### Impact of Elastic Net on classification performance

To evaluate effect of Elastic Net on the model performance, Stack Autoencoder-Kernel Density Estimation (SK) and ESK-model were compared in Figs 7-9. Evaluation method has been defined in previous section. As shown in Figs 7-9, all evaluation indicators of ESK-model were higher than SK-model that without adding Elastic Net.

**Fig 7.**
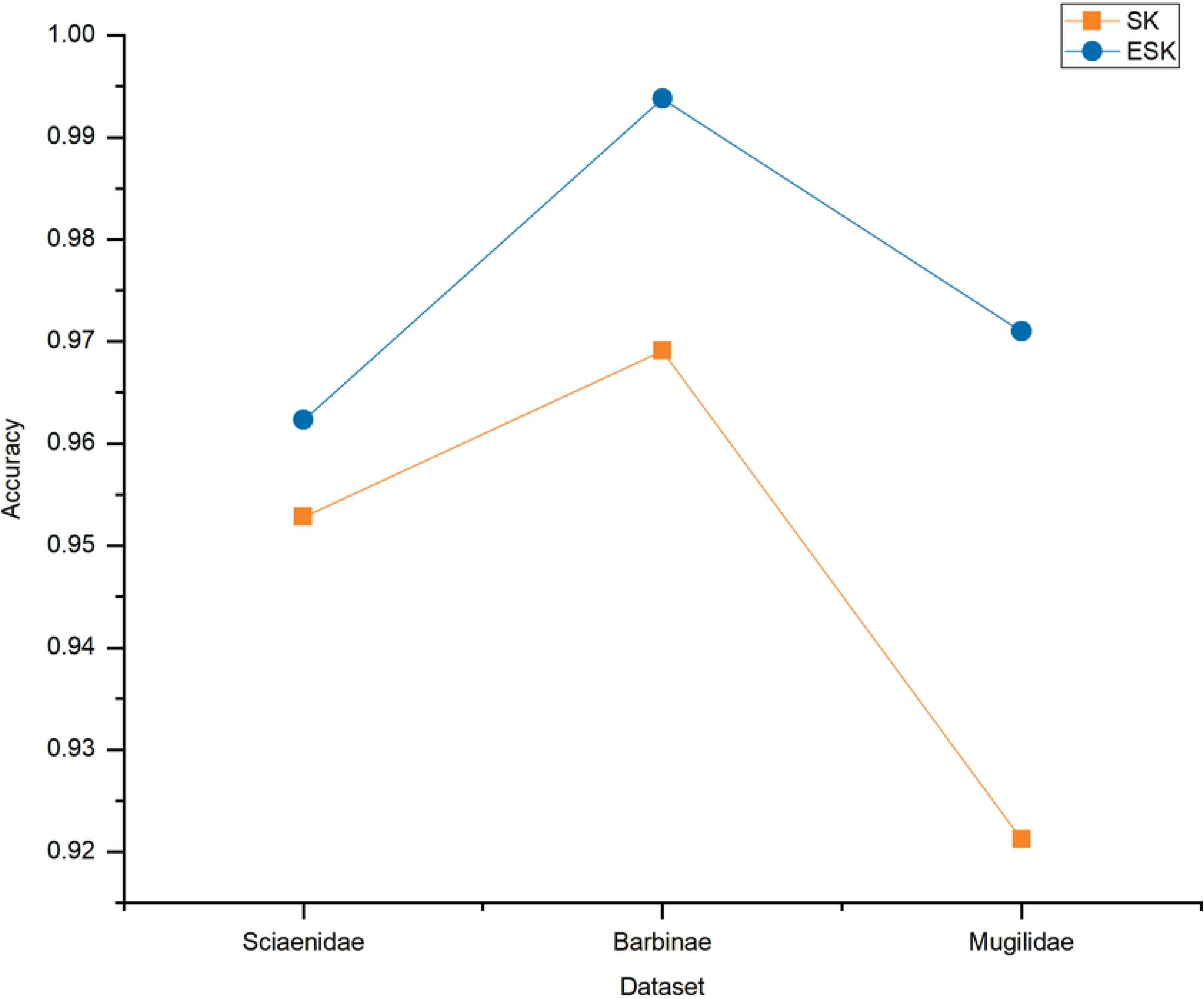
Accuracy on SK and ESK.

**Fig 8.**
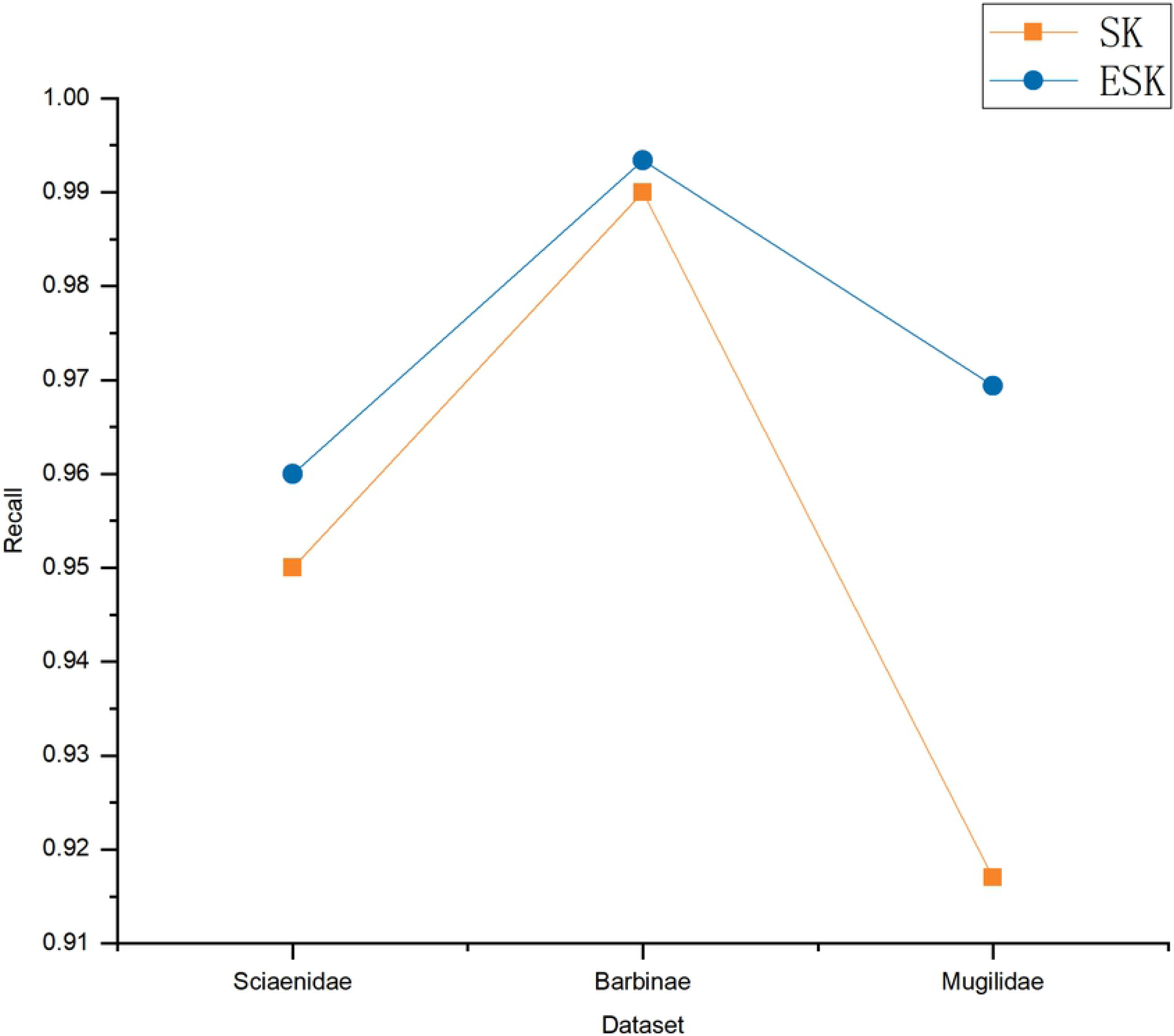
Recall on SK and ESK.

**Fig 9.**
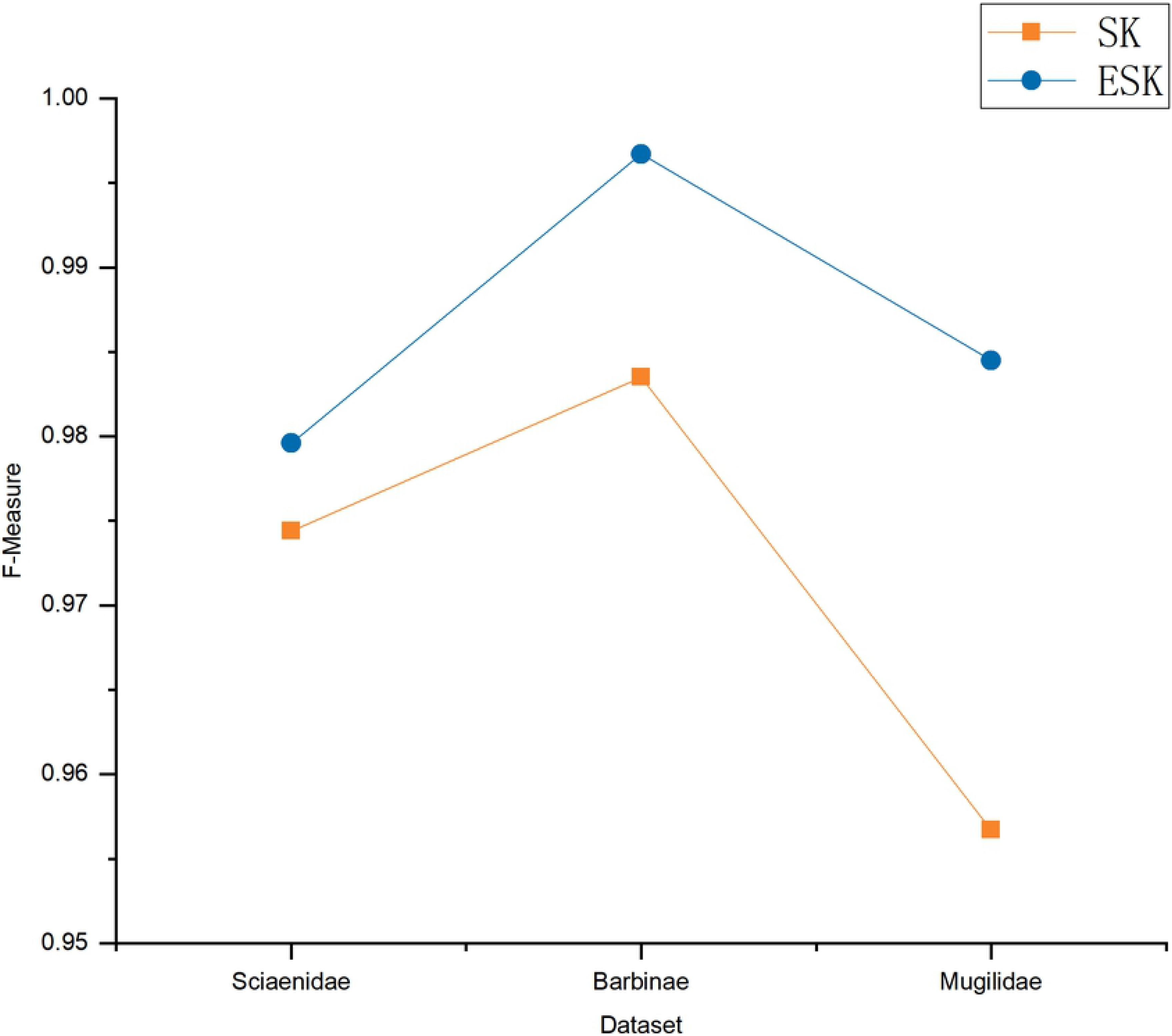
F-Measure on SK and ESK.

In addition, Table 4 illustrates the detailed data corresponding to Figs 7-9. The evaluation matrix (Accuracy, Recall, F-Measure) on Sciaenidae dataset increased by approximately 0.0095, 0.0100 and 0.0052, respectively. Similarly, under the same conditions, the evaluation matrix also increased in other two datasets. Those results indicate that add Elastic Net can improve the performance of the ESK-model.

**Table 4.** The evaluation matrix on SK and ESK models.

|  | Sciaenidae | Barbinae | Mugilidae |
| --- | --- | --- | --- |
| SK | 0.9528 0.9500 0.9744 | 0.9691 0.9900 0.9835 | 0.9212 0.9170 0.9567 |
| ESK | 0.9623 0.9600 0.9796 | 0.9938 0.9934 0.9967 | 0.9710 0.9694 0.9845 |

Note that the order of evaluation matrix is as follows Accuracy, Recall, F-measure.

### Performance evaluation with different methods

We compared our method, ESK-model, with four state-of-art algorithms, one class-support vector machine(OC-SVM) [35], K-nearest neighbor(KNN) [36], isolation Forest(iForest) [37], autoencoder(AE) [38], to evaluate performance on the task of sorting fishes from different families base on DNA barcode. Cross validation was used for model training, and confusion matrix of different models on three datasets is shown in Fig 10.

**Fig 10.**
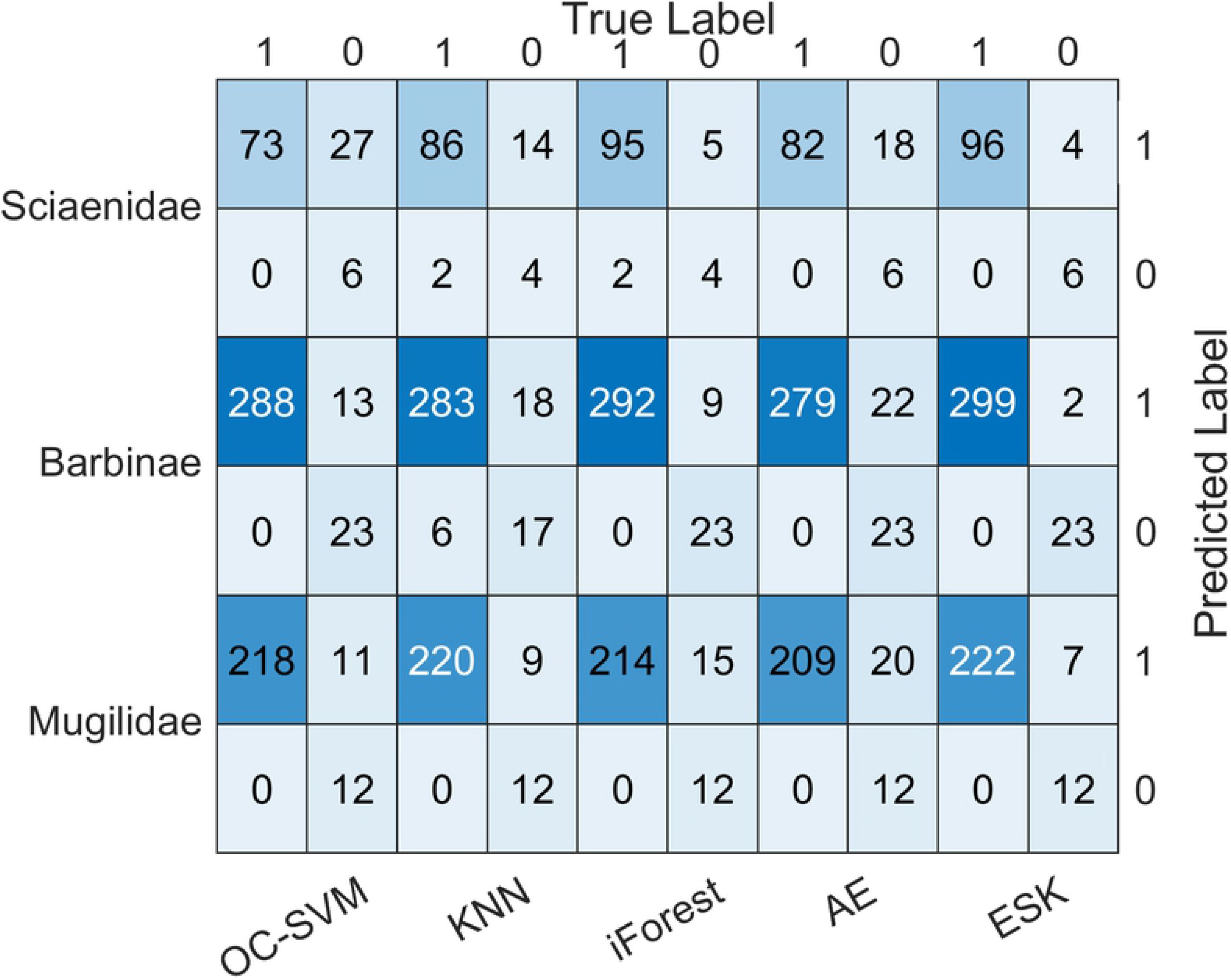
Confusion matrix of five models on three datasets.

In order to show the specific relationship between our method and other four methods, we utilize histograms to compare the performance of three matrices. Additionally, Table 5 exhibits the detailed data corresponding to Figs 11-13. As we can see in Figs 11-13, ESK-model provides stable and efficient effects on three datasets and generates the highest Accuracy, Recall and F-measure. Those results show that ESK-model is superior to other methods.

**Table 5.** The evaluation matrix of three datasets.

|  | OC-SVM | KNN | iForest | AE | ESK |
| --- | --- | --- | --- | --- | --- |
| Sciaenidae | 0.7453 0.7300 0.8439 | 0.8491 0.8600 0.9149 | 0.9340 0.9500 0.9645 | 0.8302 0.8200 0.9011 | **0.9623 0.9600 0.9796** |
| Barbinae | 0.9599 0.9568 0.9779 | 0.9228 0.9402 0.9577 | 0.9722 0.9701 0.9848 | 0.9321 0.9269 0.9621 | **0.9938 0.9934 0.9967** |
| Mugilidae | 0.9544 0.9520 0.9754 | 0.9627 0.9607 0.9800 | 0.9378 0.9345 0.9661 | 0.9170 0.9127 0.9543 | **0.9710 0.9694 0.9845** |

Note that the best result is typeset in bold. The order of evaluation matrix is as follows Accuracy, Recall, F-measure.

**Fig 11.**
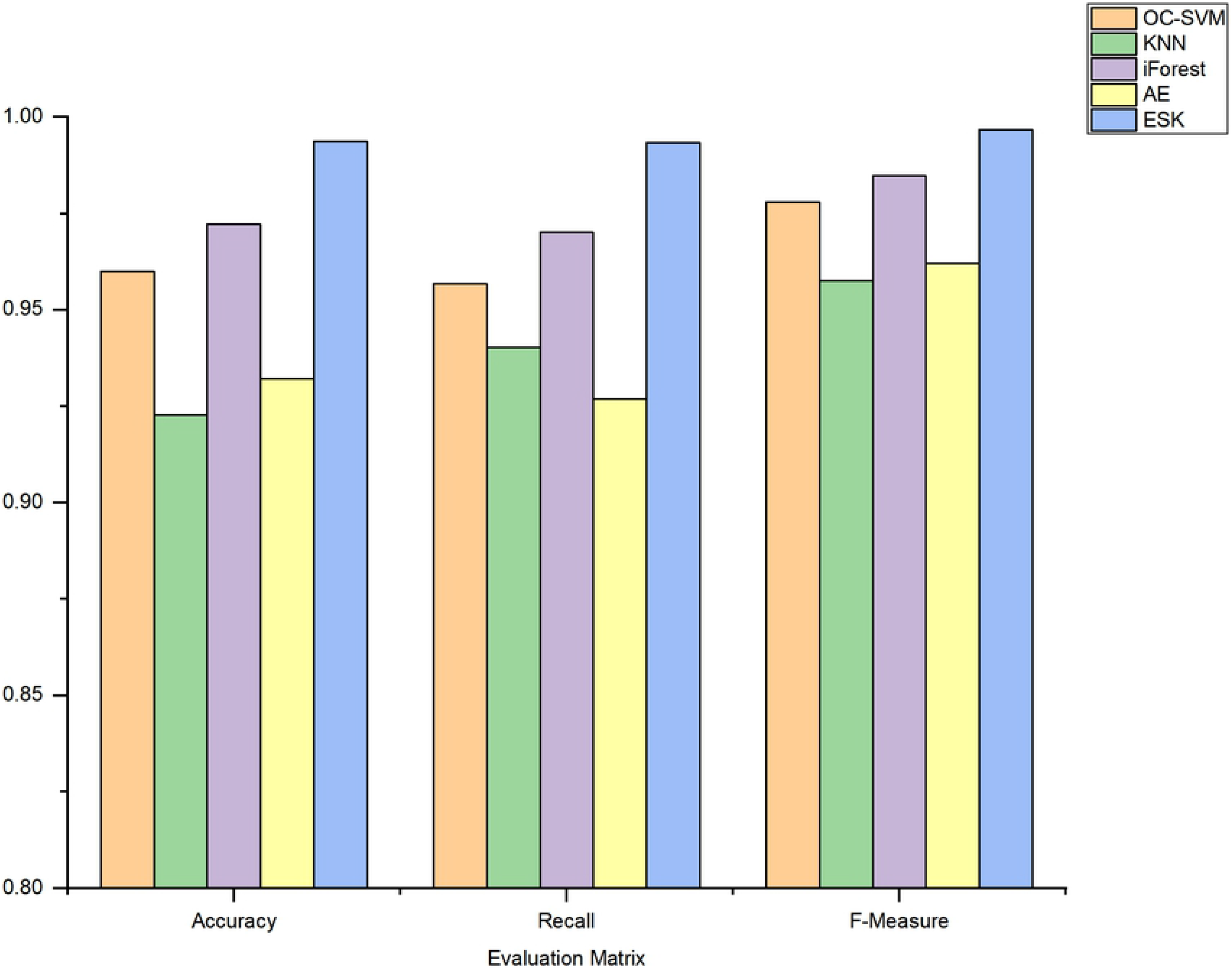
The evaluation matrix on Sciaenidae.

**Fig 12.**
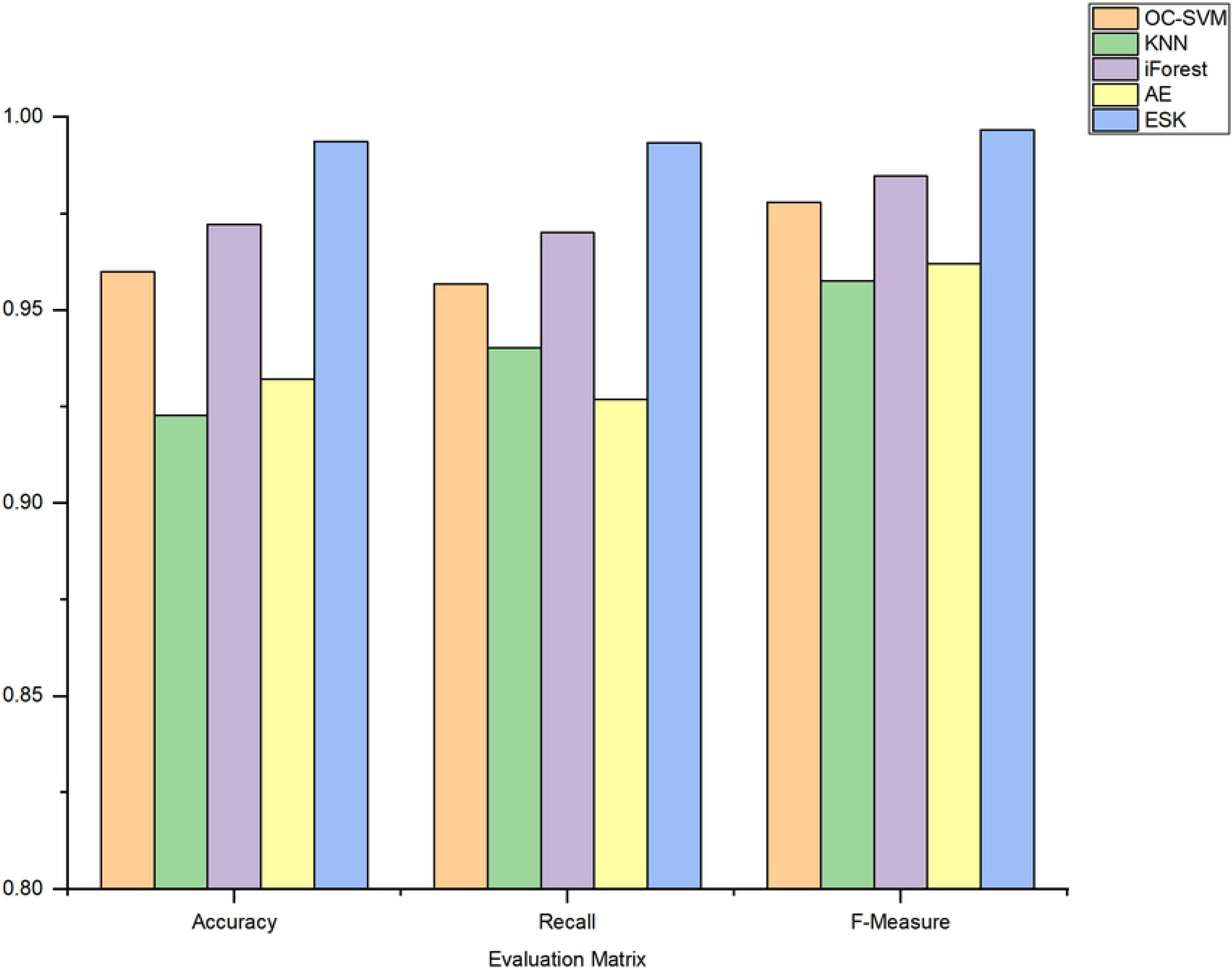
The evaluation matrix on Barbinae.

**Fig 13.**
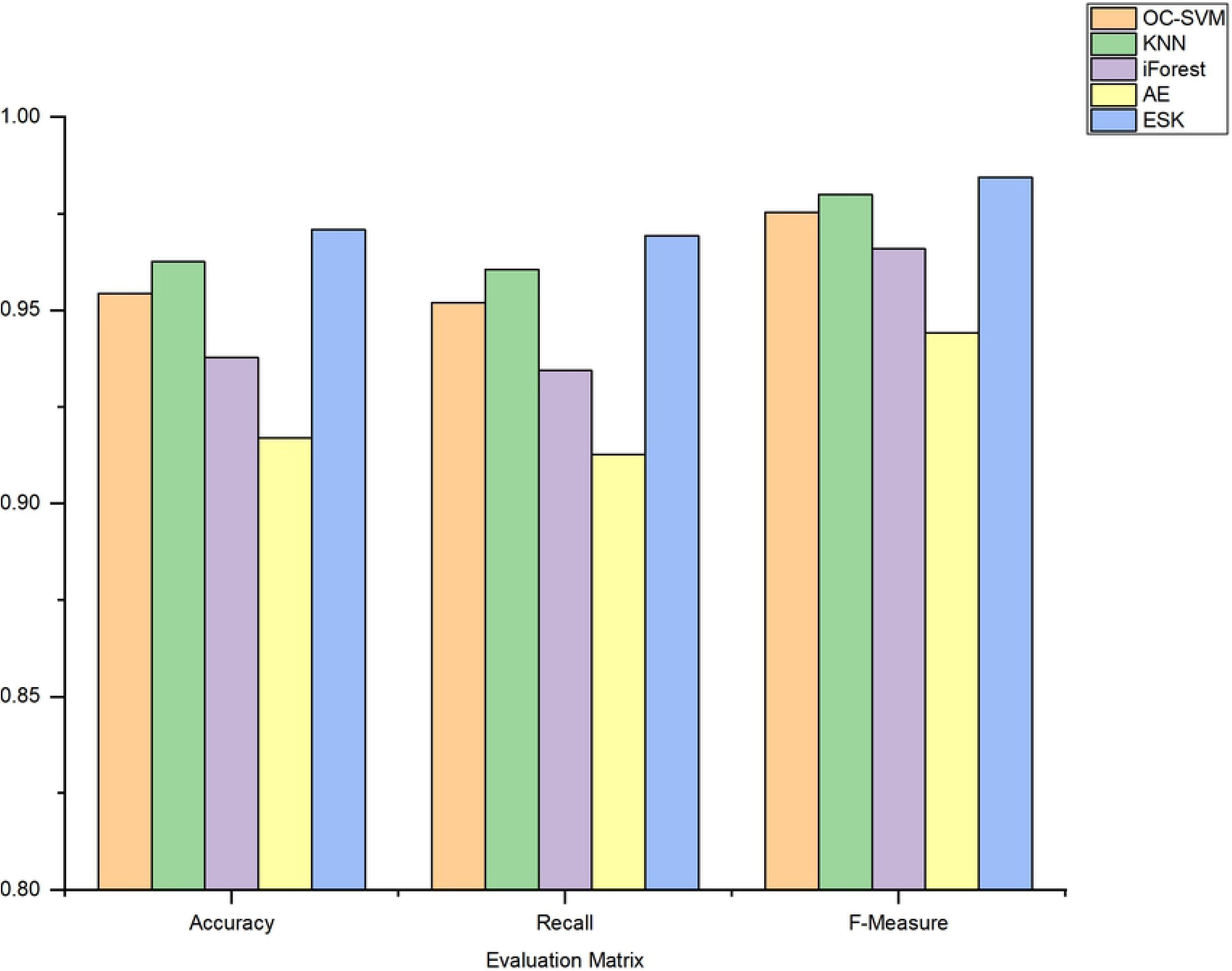
The evaluation matrix on Mugilidae.

## Discussion

This study set out with aim of constructing a novel deep learning model base on DNA barcode with the employ of representative data to classify fishes from different families and distinguish the outgroup. In this section, we discuss and analyze the experimental results and findings.

A significant experimental result was that ESK-model achieved the best discrimination performance when the number of stacked AEs was set to five. There are several possible reasons for this result. The features of COI fragment can’t be fully learned when the number of stacked AEs is few. With the increase of the number of AEs, the proposed model can learn the deeper hidden features of DNA sequences. Obviously, when the number of AEs increased to five, the outgroup scores decreased sharply. Experiments showed that increased the number of AEs did not improve performance. The performance tended to be stable when the number of AEs was more than five because the deep features had already fully learned. Hence, the prime number of stacked AEs in the ESK-model was five.

Another considerable experimental result was that Elastic Net can improve the performance of proposed model. A good model of deep learning usually requires abundant data to training, while the limitation of obtaining the COI sequences of fishes from different families, the problem of overfitting in small datasets is more and more serious. To solve the overfitting problem in training process on small datasets is of great importance. This model puts forward by using Elastic Net to solve overfitting problem and improve the generalization ability of the model. Moreover, genetic characteristics of fish belong to high-dimensional data, which is time-consuming during training. However, directly combining a set of fully connected EN-SAE is often useless to extract useful information. Elastic Net provides sparse connection also can save training time. Therefore, Elastic Net can improve the performance of proposed model.

The most surprising finding was that the proposed model could accurately classify fish from different families. EN-SAE is used to calculate the outgroup scores, when the outgroup scores are high, the probability of being identified as other families is increased. The size of fish belonging to the same family is far more than that from other families, EN-SAE can well fit and learn the characteristics of intraspecific fish in the process of training. On the contrary, the number of fishes in different families is relatively small, we can’t get a good fitting effect, resulting in higher outgroup scores. Therefore, they are more likely to be treated as outgroup in KDE-model. At the same time, compared with other algorithms, it further confirms that the proposed model has better performance in fish classification.

These positive results and findings suggest that the ESK-model based on deep learning, with the utilization of DNA barcode technology, can effectively classify the fish from different families.

## Conclusion

In this study, we proposed the ESK-model that fuses EN-SAE model and KDE technology for fish classification in different families through DNA barcode. The experimental results and findings demonstrate the effectiveness of proposed model.

The main results and findings of this paper are as follows:

1. The outgroup scores have leveled off when the number of stacked AEs was set to five.
2. Adding Elastic Net can prevent overfitting more effectively and improve the generalization ability of the model.
3. Compared with the current popular methods, our proposed model had better performance in fish classification from different families by using COI sequences.

## Supporting information

**S1 Table. Species of experimental samples on Sciaenidae.**

**S2 Table. Species of experimental samples on Barbinae.**

**S3 Table. Species of experimental samples on Mugilidae.**

## References

1. Xu L, Wang X, Van Damme K, Huang D, Li Y, Wang L, et al. Assessment of fish diversity in the South China Sea using DNA taxonomy. Fisheries Research. 2021;233:105771. doi: 10.1016/j.fishres.2020.105771.

2. Fautin D, Dalton P, Incze LS, Leong J-AC, Pautzke C, Rosenberg A, et al. An Overview of Marine Biodiversity in United States Waters. PloS one. 2010;5(8):e11914. doi: 10.1371/journal.pone.0011914.

3. Knowlton N, Weigt LA. New dates and new rates for divergence across the Isthmus of Panama. Proceedings of the Royal Society B: Biological Sciences. 1998;265(1412):2257–63. doi: 10.1098/rspb.1998.0568.

4. Thu PT, Huang WC, Chou TK, Van NQ, Liao TY. DNA barcoding of coastal ray-finned fishes in Vietnam. PloS one. 2019;14(9):e0222631. doi: 10.1371/journal.pone.0222631.

5. Hebert PD, Cywinska A, Ball SL, deWaard JR. Biological identifications through DNA barcodes. Proceedings Biological sciences. 2003;270(1512):313–21. doi: 10.1098/rspb.2002.2218. PubMed PMID: 12614582; PubMed Central PMCID: PMC1691236.

6. Ramirez JL, Rosas-Puchuri U, Canedo RM, Alfaro-Shigueto J, Ayon P, Zelada-Mazmela E, et al. DNA barcoding in the Southeast Pacific marine realm: Low coverage and geographic representation despite high diversity. PloS one. 2020;15(12):e0244323. doi: 10.1371/journal.pone.0244323. PubMed PMID: 33370342; PubMed Central PMCID: PMC7769448.

7. Liang H, Meng Y, Luo X, Li Z, Zou G. Species identification of DNA barcoding based on COI gene sequences in Bagridae catfishes. Journal of Fishery Sciences of China. 2018;25(4):772. doi: 10.3724/sp.j.1118.2018.18036.

8. Xu L, Van Damme K, Li H, Ji Y, Wang X, Du F. A molecular approach to the identification of marine fish of the Dongsha Islands (South China Sea). Fisheries Research. 2019;213:105–12. doi: 10.1016/j.fishres.2019.01.011.

9. Ren BQ, Xiang XG, Chen ZD. Species identification of Alnus (Betulaceae) using nrDNA and cpDNA genetic markers. Mol Ecol Resour. 2010;10(4):594–605. doi: 10.1111/j.1755-0998.2009.02815.x. PubMed PMID: 21565064.

10. Newmaster SG, Fazekas AJ, Steeves RAD, Janovec J. Testing candidate plant barcode regions in the Myristicaceae. Molecular Ecology Resources. 2008;8(3):480–90. doi: 10.1111/j.1471-8286.2007.02002.x.

11. Liu J, Moller M, Gao LM, Zhang DQ, Li DZ. DNA barcoding for the discrimination of Eurasian yews (Taxus L., Taxaceae) and the discovery of cryptic species. Mol Ecol Resour. 2011;11(1):89–100. doi: 10.1111/j.1755-0998.2010.02907.x. PubMed PMID: 21429104.

12. Necchi O, West JA, Rai SK, Ganesan EK, Rossignolo NL, de Goër SL. Phylogeny and morphology of the freshwater red algaNemalionopsis shawii(Rhodophyta, Thoreales) from Nepal. Phycological Research. 2016;64(1):11–8. doi: 10.1111/pre.12116.

13. Valentini A, Pompanon F, Taberlet P. DNA barcoding for ecologists. Trends in ecology & evolution. 2009;24(2):110–7. doi: 10.1016/j.tree.2008.09.011. PubMed PMID: 19100655.

14. Ji Y, Ashton L, Pedley SM, Edwards DP, Tang Y, Nakamura A, et al. Reliable, verifiable and efficient monitoring of biodiversity via metabarcoding. Ecology letters. 2013;16(10):1245–57. doi: 10.1111/ele.12162. PubMed PMID: 23910579.

15. Gathier G, van der Niet T, Peelen T, van Vugt RR, Eurlings MC, Gravendeel B. Forensic identification of CITES protected slimming cactus (Hoodia) using DNA barcoding. Journal of forensic sciences. 2013;58(6):1467–71. doi: 10.1111/1556-4029.12184. PubMed PMID: 23865560.

16. Liu J, Yan H-F, Newmaster SG, Pei N, Ragupathy S, Ge X-J, et al. The use of DNA barcoding as a tool for the conservation biogeography of subtropical forests in China. Diversity and Distributions. 2015;21(2):188–99. doi: 10.1111/ddi.12276.

17. Wang T, Qi D, Sun S, Liu Z, Du Y, Guo S, et al. DNA barcodes and their characteristic diagnostic sites analysis of Schizothoracinae fishes in Qinghai province. Mitochondrial DNA Part A. 2019; 30(4):592–601. doi: 10.1080/24701394.2019.1580273. Epub 2019 Apr 5. PMID: 30952197.

18. Hebert PDN, Stoeckle MY, Zemlak TS, Francis CM. Identification of Birds through DNA Barcodes. PLOS Biology. 2004;2(10):e312. doi: 10.1371/journal.pbio.0020312.

19. Kerr KCR, Stoeckle MY, Dove CJ, Weigt LA, Francis CM, Hebert PDN. Comprehensive DNA barcode coverage of North American birds. Molecular Ecology Notes. 2007; 7(4):535–543. doi: 10.1111/j.1471-8286.2007.01670.x. PMID: 18784793; PMCID: PMC2259444.

20. Gang, Wang, Chunxiao, Li, Xiaoxia, Guo, et al. Identifying the Main Mosquito Species in China Based on DNA Barcoding. PloS one. 2012; 7(10): e47051. https://doi.org/10.1371/journal.pone.0047051.

21. Zhang J. Species identification of marine fishes in china with DNA barcoding. Evidence-based complementary and alternative medicine: eCAM. 2011; 8(1):1–10. doi: 10.1155/2011/978253. PubMed PMID: 21687792; PubMed Central PMCID: PMC3108176.

22. Steinke D, Zemlak TS, Boutillier JA, Hebert PDN. DNA barcoding of Pacific Canada’s fishes. Marine Biology. 2009;156(12):2641–7. doi: 10.1007/s00227-009-1284-0.

23. Thu PT, Huang WC, Chou TK, Van Quan N, Van Chien P, Li F, et al. DNA barcoding of coastal ray-finned fishes in Vietnam. PloS one. 2019;14(9):e0222631. doi: 10.1371/journal.pone.0222631. PubMed PMID: 31536551; PubMed Central PMCID: PMC6752846.

24. Talaga S, Leroy C, Guidez A, Dusfour I, Girod R, Dejean A, et al. DNA reference libraries of French Guianese mosquitoes for barcoding and metabarcoding. PloS one. 2017;12(6):e0176993. doi: 10.1371/journal.pone.0176993. PubMed PMID: 28575090; PubMed Central PMCID: PMC5456030.

25. Decru E, Moelants T, De Gelas K, Vreven E, Verheyen E, Snoeks J. Taxonomic challenges in freshwater fishes: a mismatch between morphology and DNA barcoding in fish of the north-eastern part of the Congo basin. Mol Ecol Resour. 2016;16(1):342–52. doi: 10.1111/1755-0998.12445. PubMed PMID: 26186077.

26. Iyiola OA, Nneji LM, Mustapha MK, Nzeh CG, Oladipo SO, Nneji IC, et al. DNA barcoding of economically important freshwater fish species from north-central Nigeria uncovers cryptic diversity. Ecology and evolution. 2018;8(14):6932–51. doi: 10.1002/ece3.4210. PubMed PMID: 30073057; PubMed Central PMCID: PMC6065348.

27. Wang T, Qi D, Sun S, Liu Z, Du Y, Guo S, et al. DNA barcodes and their characteristic diagnostic sites analysis of Schizothoracinae fishes in Qinghai province. Mitochondrial DNA Part A, DNA mapping, sequencing, and analysis. 2019;30(4):592–601. doi: 10.1080/24701394.2019.1580273. PubMed PMID: 30952197.

28. Ward RD, Hanner R, Hebert PD. The campaign to DNA barcode all fishes, FISH-BOL. Journal of fish biology. 2009;74(2):329–56. doi: 10.1111/j.1095-8649.2008.02080.x. PubMed PMID: 20735564.

29. Jin S, Zeng X, Xia F, Huang W, Liu X. Application of deep learning methods in biological networks. Briefings in bioinformatics. 2020;bbaa043. doi: 10.1093/bib/bbaa043. PubMed PMID: 32363401.

30. Chu Z, Yu J. An end-to-end model for rice yield prediction using deep learning fusion. Computers and Electronics in Agriculture. 2020;174:105471. doi: 10.1016/j.compag.2020.105471.

31. Chen J, Sathe S, Aggarwal C, Turaga D. Outlier Detection with Autoencoder Ensembles. Proceedings of the 2017 SIAM International Conference on Data Mining (SDM). 2017;pp. 90–98. dio: 10.1137/1.9781611974973.11.

32. Homoliak I. Convergence Optimization of Backpropagation Artificial Neural Network Used for Dichotomous Classification of Intrusion Detection Dataset. Journal of Computers. 2017;143–55. doi: 10.17706/jcp.12.2.143-155.

33. Vincent P, Larochelle H, Lajoie I, Bengio Y, Manzagol PA. Stacked Denoising Autoencoders: Learning Useful Representations in a Deep Network with a Local Denoising Criterion. Journal of Machine Learning Research. 2010;11(12):3371–408.

34. Taaffe K, Pearce B, Ritchie G. Using kernel density estimation to model surgical procedure duration. International Transactions in Operational Research. 2018;28(1):401–18. doi: 10.1111/itor.12561.

35. Erfani SM, Rajasegarar S, Karunasekera S, Leckie C. High-dimensional and large-scale anomaly detection using a linear one-class SVM with deep learning. Pattern Recognition. 2016;58:121–34. doi: 10.1016/j.patcog.2016.03.028.

36. Hastie T, Tibshirani R. Discriminant adaptive nearest neighbor classification. IEEE Transactions on Pattern Analysis & Machine Intelligence. 1996;18(6):607–16.

37. Liu FT, Ting KM, Zhou ZH. Isolation-Based Anomaly Detection. Acm Transactions on Knowledge Discovery from Data. 2012;6(1):1–39.

38. Guo J, Liu G, Zuo Y, Wu J, editors. An Anomaly Detection Framework Based on Autoencoder and Nearest Neighbor. 2018 15th International Conference on Service Systems and Service Management (ICSSSM); 2018;pp. 1–6, doi: 10.1109/ICSSSM.2018.8464983.

